# Posterior Cortex Isolation Enhances Detection of Alpha Desynchronization During Sustained Attention

**DOI:** 10.1101/2025.11.17.688558

**Authors:** N St. Clair, S Mahajan, C Leung, C Srinivas, L Oushana, N Dewan, F Zhou

## Abstract

Electroencephalography (EEG) is an essential tool for assessing the neural activity behind mental fatigue, a factor that can influence both cognitive and physical performance. Mental fatigue influences alpha-band oscillations (8–12 Hz), reflecting cortical engagement and attention stability. In this study, EEG recordings from 109 participants in the Physionet open-source EEG Motor Movement/Imagery Dataset (EEGMMIDB) were analyzed to characterize alpha power changes during sustained attention. Spectral analysis was performed across early and late task blocks to quantify alpha desynchronization over time. Participants demonstrated a consistent decrease in alpha power (Δ = −0.076 ± 0.019 log_10_ units; *p* = 0.0001), indicating desynchronization associated with sustained attentional demand. These findings show the applicability of open-source EEG datasets for research. Furthermore, these findings support the isolation of posterior activation as a marker of mental fatigue through alpha desynchronization.

**One sentence overview:** Alpha desynchronization is a marker for mental fatigue in EEGs. This paper illustrates that isolating the posterior cortical lobes enhances the resolution and sensitivity of alpha band changes associated with sustained attention.

## Introduction

Among the various EEG frequency ranges the alpha frequency has the most evident disengagement in synchronized cortical activation. In response to stimulus the alpha range experiences an increase or decrease in amplitude depending on the synchronization or desynchronization of the rhythmic neuropotentials. These neuropotentials are driven through thalamo-cortical loops which connect the thalamus to the posterior cortex driving the formation of alpha waves. The alpha frequency captures resting cortical activation due to synchronized, rhythmic firing of neurons (Benedek et al., 2014; Klimesch, 1999; Klimesch, 2012; Mathewson et al., 2009). During sustained attention rhythmic activation of neurons desynchronizes and the alpha power decreases (Tanaka et al., 2012; Clayton et al., 2015). While the role of alpha waves is established, the involvement in the posterior cortex in desynchronization of these waves is more abstract. Isolation of the posterior cortex when examining alpha wave fluctuations over time may provide greater resolution and reproducibility. It is expected that the alpha power decreases between the early and late segments of a sustained attention task, reflecting neural desynchronization associated with mental fatigue. Upon isolation of the posterior cortex this alpha desynchronization should be greater and better resolved (Boksem, Meijman & Lorist, 2005; Boksem & Tops, 2008; Liu et al., 2019).

## Methods

### Physionet Database

The sample size for this study included 109 participants that were supplied from the PhysioBank database which recorded 64-channel EEG results for a variety of simple motor and visualization tasks (Goldberger et al., 2000). Each participant was selected on a voluntary basis, and were all healthy adults. The PhysioBank database collected multiple samples of one to two minute EEG recordings for each participant, and gathered a total of 1500 recordings.

This study was conducted as a within-subjects design, as each participant was tasked with participating in 6 different tasks and after each task, the participants were instructed to relax. The study design had participants complete both of the baseline runs first, starting with keeping one’s eyes open, and then complete tasks three, four, five, and six in that order a total of three times. However, only the eyes-closed resting-state condition was analyzed in the present study. This condition provides a stable resting baseline with robust posterior alpha activity and minimal motor or cognitive confounds, making it optimal for detecting desynchronization related to sustained attention and mental fatigue (Mathewson et al., 2009). Each eyes-closed run lasted approximately one minute, yielding 109 individual recordings suitable for group-level spectral analysis.

### EEG Processing

All data were processed using the Brainwave EEG OpenLab framework (Python 3.11, MNE-Python v1.6.0). Raw.edf files were retrieved from the PhysioNet repository, organized under a standardized directory structure (data/raw), and all derived products were saved in (data/derivatives). Continuous EEG signals were band-pass filtered between 0.5 and 30 Hz to remove slow drifts and high-frequency noise, and a 60 Hz notch filter was applied to suppress line interference. Signals were re-referenced to the common average across all 64 scalp electrodes. All 109 recordings met quality-control criteria and were retained for analysis (Gramfort et al., 2014).

Each filtered recording was segmented into non-overlapping 2 s epochs. Power spectral density (PSD) was computed for every epoch using Welch’s method with a Hanning window (Welch, 1967). Alpha-band power (8–12 Hz) was extracted from the PSD and averaged across posterior electrodes (P–, PO–, O–, Iz–) to quantify occipital–parietal oscillatory activity. To examine temporal change within the one-minute recording, the first ten epochs were defined as the early block and the final ten as the late block. Alpha power values were log_10_-transformed for normalization, and paired-sample t-tests (scipy.stats.ttest_rel) were conducted comparing late < early alpha power across subjects. Group-level statistics, including mean Δ (log α power), p-values, and effect sizes, were compiled automatically in a summary table.

## Results

Across 109 participants, alpha-band (8–12 Hz) power decreased from the early to late portions of the eyes-closed resting-state recording. Topographic analyses revealed a decrease in alpha-band power across the early to late block localized primarily to posterior electrodes (P–, PO–, O–, Iz; Figure 3). Paired-sample testing confirmed a significant within-subject reduction in alpha power (Δ = –0.076 ± 0.019 log_10_ units; p = 0.0001; Figure 6). Individual subject trajectories further showed that the majority of participants displayed a negative shift in posterior alpha power over time (Figures 4 and 6). Posterior isolation produced a stronger effect size and lower p-value relative to whole-brain averaging, demonstrating improved sensitivity (Δ = –0.052 vs –0.076; Figures 4 and 6). Together, these results indicate that isolating posterior cortical regions enhances the resolution and detection of alpha desynchronization associated with sustained attention and early mental fatigue.

**Figure 1.**
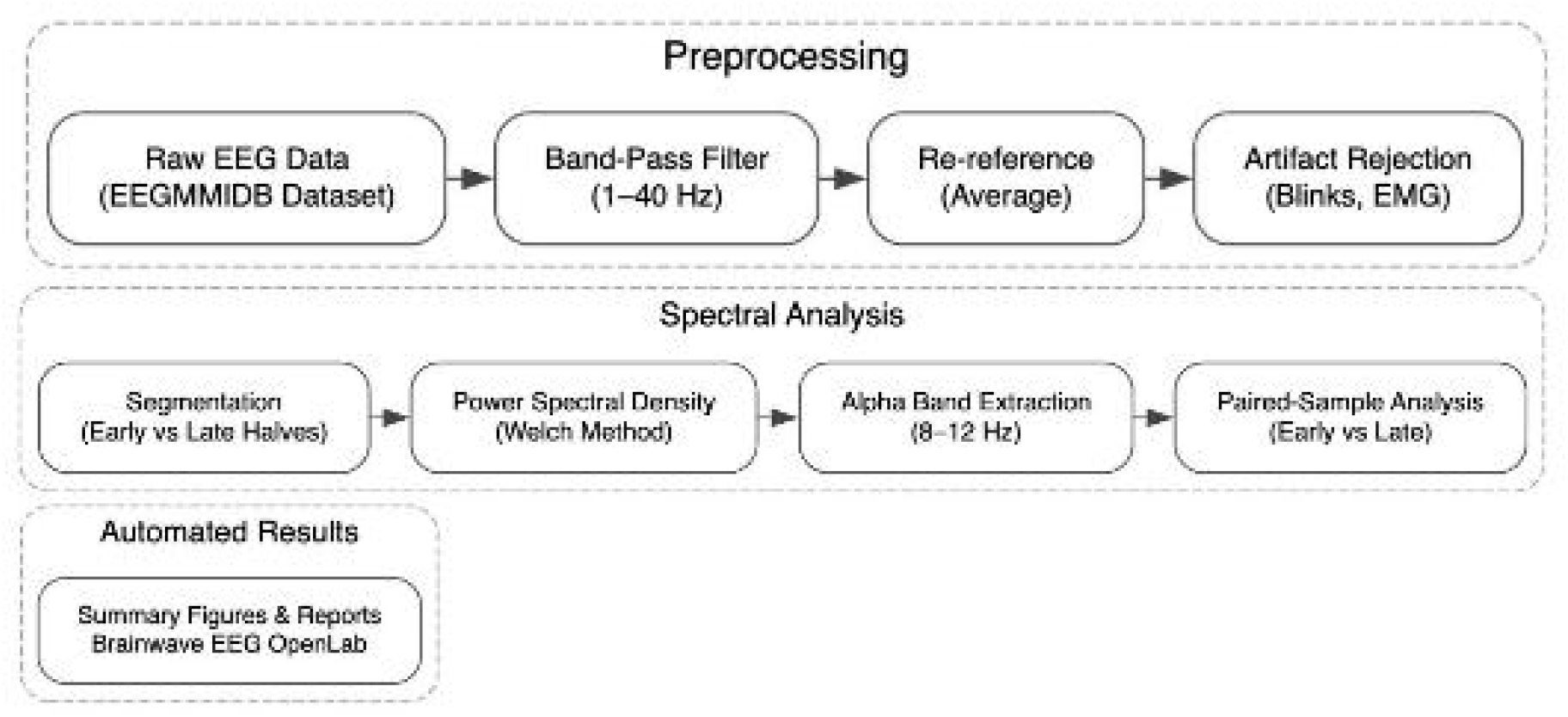
Overview of the Brainwave EEG OpenLab preprocessing and analysis pipeline. Raw EEG data from the open-source EEGMMIDB dataset were band-pass filtered (1–40 Hz), re-referenced, and cleaned for artifacts. Data were segmented into early and late halves, and power spectral density (PSD) was computed using the Welch method. Alpha-band (8–12 Hz) power was compared between blocks using a paired-sample analysis. Results were compiled into automated summary figures and reports.

**Figure 2.**
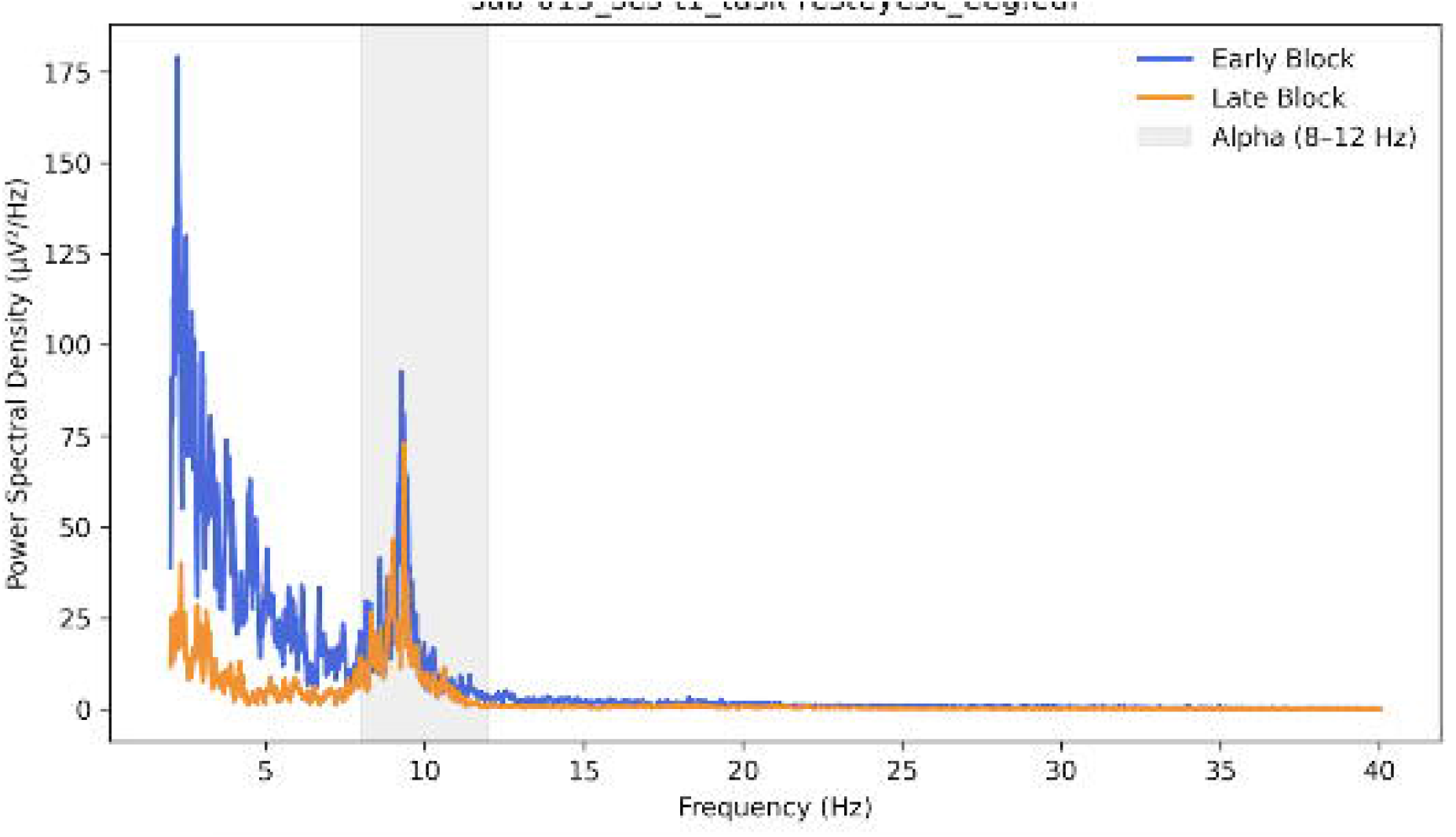
Representative power spectral density (PSD) for Subject S015R02. EEG power spectra were computed for early (blue) and late (orange) halves of the recording. The shaded region marks the alpha band (8–12 Hz), with arrows indicating the alpha peak (~10 Hz). A relative reduction in alpha-band power during the late block illustrates alpha desynchronization, consistent with increased cortical activation during sustained attention.

**Figure 3.**
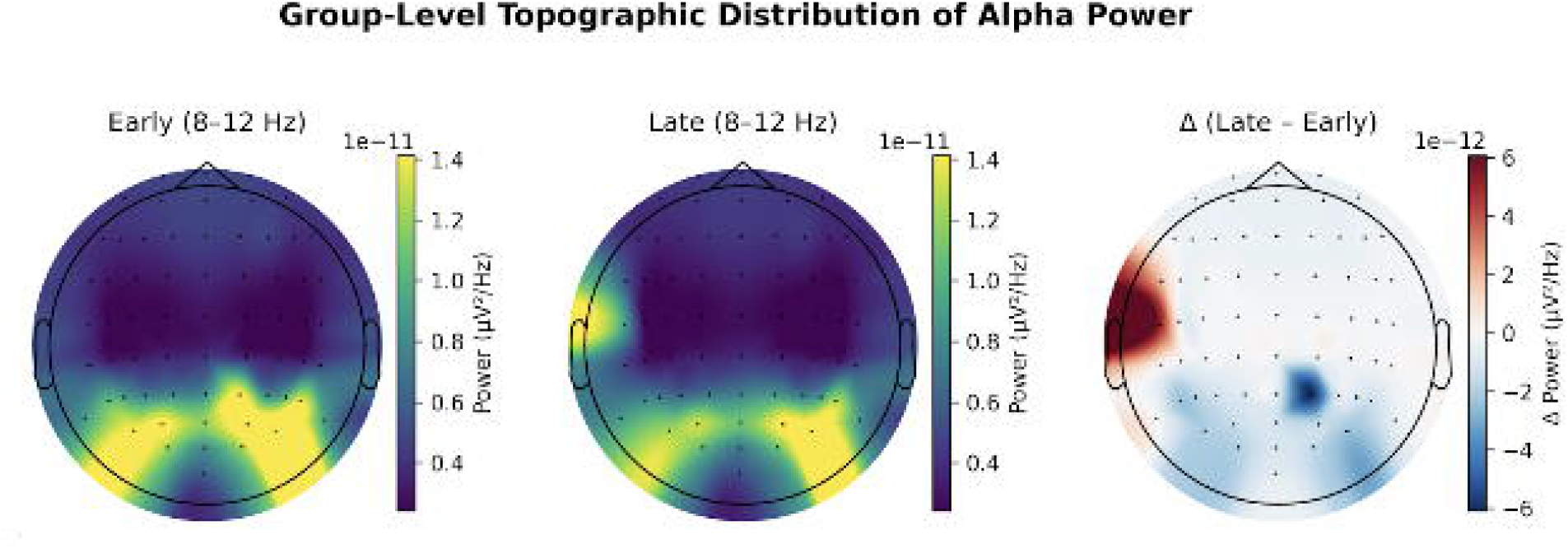
Group-level topography of alpha power during early and late blocks. Scalp maps display mean alpha-band (8–12 Hz) power averaged across all participants for the early (left) and late (middle) halves of the resting-state recording, with a difference map (right; Δ = Late – Early) illustrating relative change over time. White dots indicate posterior electrodes (P-, PO-, O-, Iz sites) included in the analysis. Blue regions on the difference map represent decreased alpha power in the late block, consistent with posterior alpha desynchronization—a neural signature of increasing mental fatigue during sustained rest.

**Figure 4.**
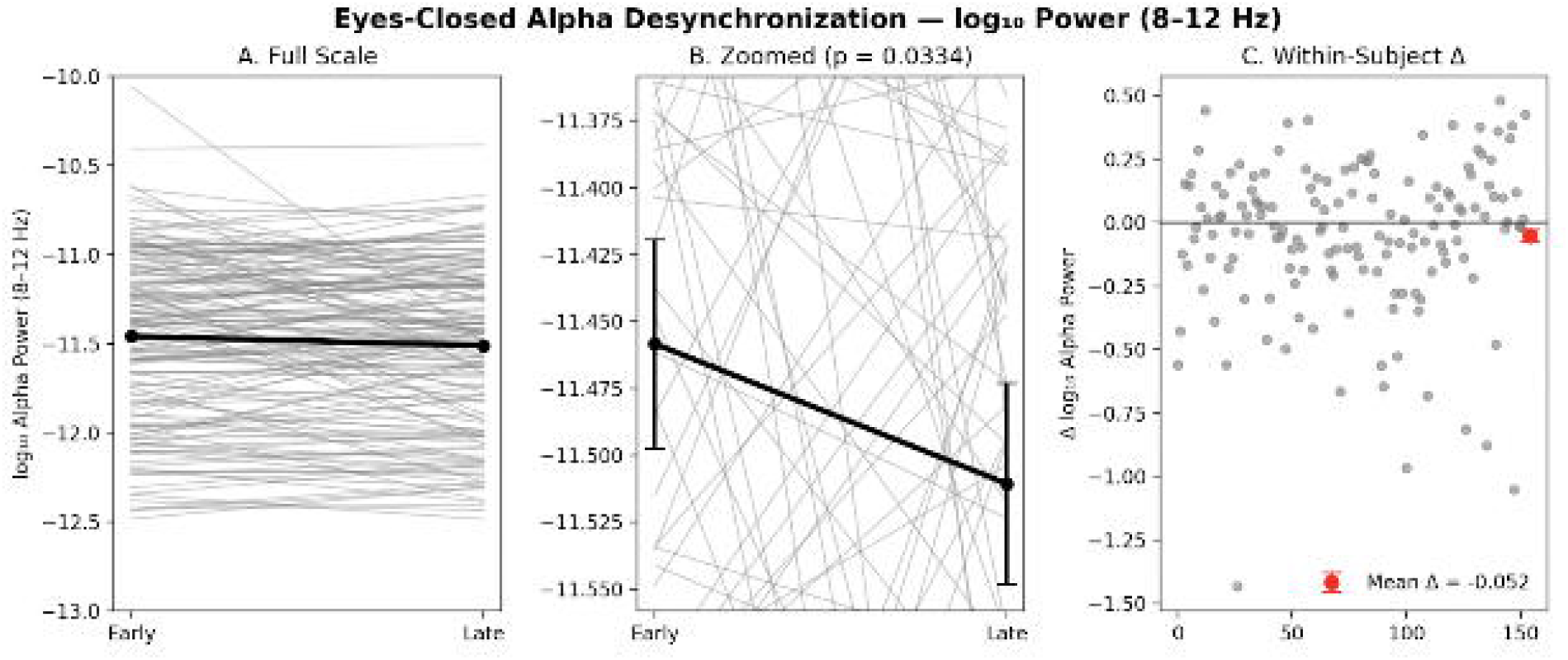
Eyes-closed alpha desynchronization over time during baseline rest. Panel A shows absolute log-transformed alpha power (8–12 Hz) across all subjects during early versus late segments of the one-minute eyes-closed baseline. Each gray line represents an individual subject, and the thick black line indicates the group mean. Panel B presents the same data on a zoomed scale, illustrating a small but statistically significant decrease in alpha power across time (paired *t*-test, *p* = 0.0334). Panel C displays within-subject changes in log_10_ alpha power (Late – Early), with the red marker denoting the mean ± SEM (Δ = −0.052 log_10_ units). Together, these results indicate a modest alpha desynchronization—consistent with reduced cortical idling or increased alertness—over the course of the eyes-closed baseline period.

**Figure 5.**
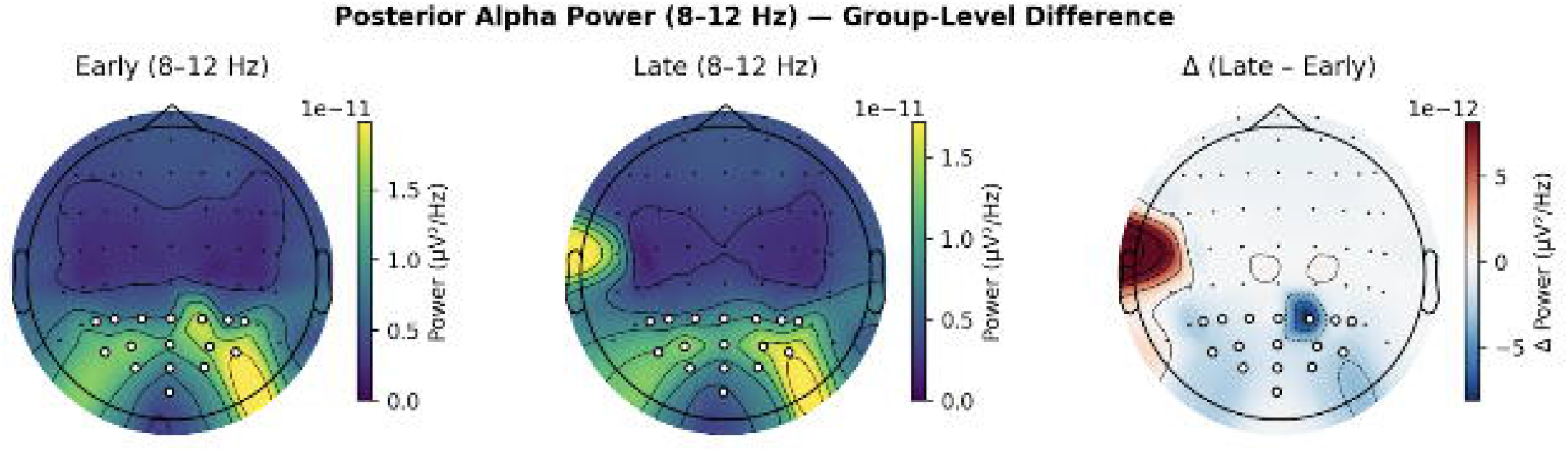
Posterior Alpha Power (8–12 Hz) — Group-Level Difference. Topographic scalp maps show mean alpha-band (8–12 Hz) power across all subjects for the early (left) and late (middle) halves of the eyes-closed resting session, and their difference (right, Δ = Late – Early). White dots mark the posterior electrodes (P–, PO–, O–, Iz) included in the analysis. Blue regions in the difference map indicate a reduction in posterior alpha power over time, reflecting alpha desynchronization consistent with increasing mental fatigue during sustained rest.

**Figure 6.**
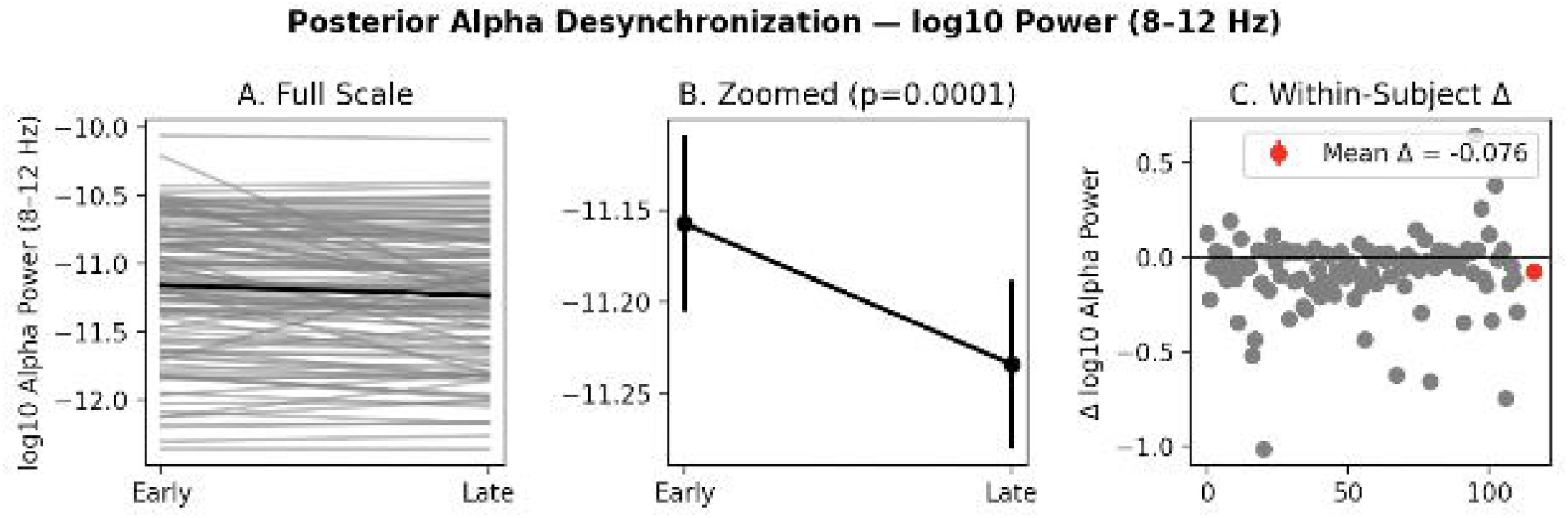
Posterior alpha desynchronization across subjects during sustained rest. Panels show within-subject changes in log_10_-transformed alpha power (8–12 Hz) between the early and late halves of the session, averaged across posterior electrodes (P–, PO–, O–, Iz). (A) Individual trajectories with mean trend line overlaid. (B) Zoomed mean ± SEM showing a significant decrease in posterior alpha power (paired *t*(109) = –3.97, *p* = 0.0001). (C) Distribution of within-subject differences (Δ log_10_ α power), with a group mean decrease of –0.076 ± 0.019. This reduction in posterior alpha power indicates significant alpha desynchronization, consistent with neural markers of mental fatigue during sustained attention.

## Discussion

This study shows that alpha desynchronization indicates an increase in cortical activation under sustained attention load. Alpha oscillations typically range from 8-12 Hz, and are usually prominent during relaxed wakefulness and are thought to reflect an “idling” state of cortical networks. When attentional load increases, however, cortical regions become more actively engaged in processing task-relevant information. This heightened activation disrupts the firing pattern characteristics of the alpha rhythm, resulting in decreased alpha power. Such desynchronization has been consistently interpreted as an electrophysiological marker of increased neural engagement and cognitive effort. Therefore, the findings support the notion that reduced alpha amplitude reflects a shift from a resting, low-demand neural state to one characterized by active sensory processing and sustained attention (Benedek et al., 2014; Provenza et al., 2019; Klimesch, 1999).

These results align with prior studies demonstrating that alpha desynchronization accompanies sustained attention and increased cognitive demand (Klimesch, 1999; Clayton, Yeung, & Cohen Kadosh, 2015). Reductions in posterior alpha power have been shown to reflect enhanced sensory cortical excitability and top-down attentional control (Mathewson et al., 2014; Sadaghiani et al., 2012). Moreover, research on mental fatigue indicates that prolonged cognitive effort produces complex alpha dynamics—often characterized by overall decreases in alpha power as the brain maintains elevated cortical activation despite declining efficiency (Boksem, Meijman, & Lorist, 2005). Collectively, these findings support the interpretation that alpha desynchronization serves as a reliable neural marker of sustained attention and cognitive effort under extended task conditions.

Upon isolation of the posterior cortex the resolution, measured through the reduction in the p value across the paired t test Δ-0.0333, drastically improved. This result aligns with the idea that thalomo-posterior cortical connections drive the formation and sustainment of alpha waves (Liu et al., 2019). Furthermore the magnitude of alpha wave desynchronization varied substantially through posterior isolation decreasing by −0.024 log units (Liu et al., 2019; Klimesch, 2012).

## Limitations

Although the dataset with 109 participants had sufficient statistical power, expanding the sample size could have improved the robustness of the analysis. Also, our study used a simplified segmentation of blocks into early vs late, so we could have segmented more windows to obtain higher temporal resolution. Alpha desynchronization was used as a proxy for mental fatigue, but there were limited psychometric measures to assess fatigue (e.g. a visual analog scale ranging from 0-100 mm) (Brietzke et al., 2021).

Since the EEGMMIDB dataset was intended for motor and imagery instead of mental fatigue, there is the risk of confounding brain regions. Our posterior isolation excludes the frontal and central cortices, thus excluding the centrally located motor cortex. Also, we only analyzed the eyes-closed resting state, which reduces interference from task-activated regions. However, closing eyes can cause spontaneous visual imagination, which can inadvertently influence alpha power in the occipital lobe and thus posterior cortex (Wei, 2018; Zapała, 2023).

## 5.3 Future Directions

Literature reports a decrease in beta/theta ratio as correlated to mental fatigue, as well as shifts in frontal-parietal and frontal-central networks (Qi 2020). These markers could be analyzed in the EEGMMIDB data or future datasets (Raufi, 2022).

Since this study focused on resting state data, future studies can apply the same statistical pipeline to task based datasets. The Psychomotor Vigilance Task is considered a gold standard in fatigue assessment (Zhu 2024) and would be a desirable candidate for a future dataset. Other tasks known to induce mental fatigue are the stroop, n-back, and isometric handgrip task (Knoop 2023). These tasks give behavioral measures which could be used to assess fatigue across time (e.g. reaction time, accuracy, grip intensity).

This study was based on the EEGMMIDB dataset; a major future objective is to apply this methodology to original EEG data that Brainwave would collect at UCLA. This would involve collecting data from the eyes-closed resting state in order to validate our results, but could include subjective ratings and fatigue-inducing tasks.

## 6. Conclusion

This study examined alpha-band power (8-12 Hz) dynamics during sustained attention using an open-source EEG dataset and a reproducible preprocessing pipeline. Across 109 participants, a significant decrease in posterior alpha-band power was observed between the early and late phases of recording, indicating alpha desynchronization associated with increased cortical engagement and the onset of mental fatigue (Clayton et al., 2015; Klimesch, 2012). Alpha desynchronization, defined as a decrease in alpha-band (8-12 Hz) power, reflects a reduction in (synchronous oscillatory activity) within cortical networks and is commonly associated with attentional engagement (Clayton et al., 2015; Klimesch, 2012). In this study, participants’ EEG recordings during an eyes-closed resting session showed a measurable decrease in alpha power over time (Figures 4, 6), demonstrating that sustained attention during resting state EEG is sufficient to produce detectable neural charges. Quantitatively, posterior alpha power decreased by −0.076 ± 0.019 log_10_ units (paired t(111) = −3.97, p = 0.0001; Figure 6C). Topographic analyses indicated that this desynchronization was concentrated over posterior electrodes (P–, PO–, O–, Iz–; Figures 3, and 6), and individual trajectories confirmed that this trend was consistent across participants (Figures 4 and 6A-B). These findings ultimately confirm that alpha desynchronization reflects neural markers of mental fatigue during sustained attention (Boksem, Meijman, & Lorist, 2005; Tanaka et al., 2012). Importantly, isolating posterior electrodes enhanced the sensitivity of detecting these changes, reinforcing the posterior cortex’s primary role in generating and modulating alpha oscillations.

Furthermore, the results validate both the BrainWave EEG OpenLab pipeline and the EEGMMIDB open source datasets as reliable tools for EEG research. Together, the results show that sustained attention leads to measurable alpha desynchronization consistent with mental fatigue, and that isolating the posterior cortex significantly improves the detection of these neural signatures. These findings support the continued use of open-access EEG analysis pipelines and provide a foundation for future investigations into neural fatigue, attentional dynamics, and task-based EEG models (Boksem et al., 2005; Tanaka et al., 2012).

## Acknowledgments

Department of Chemistry and Biochemistry, University of California, Los Angeles, CA, 90095, USA, Noah St. Clair

Department of Neuroscience, University of California, Los Angeles, CA, 90095, USA, Nathan Dewan

Department of Physiological Sciences, University of California, Los Angeles, CA, 90095, USA, Luka Oushana

Department of Psychobiology, University of California, Los Angeles, CA, 90095, USA, Chris Leung

Department of Microbiology Immunology & Molecular Genetics, University of California, Los Angeles, CA, 90095, USA, Shaurya Mahajan

Department of Psychobiology, University of California, Los Angeles, CA, 90095, USA, Faith Zhou

